# An Eocene Origin of Passerine Birds Estimated Using Bayesian Tip Dating with Fossil Occurrences

**DOI:** 10.1101/2025.04.28.650918

**Authors:** Arong Luo, Jacqueline M.T. Nguyen, Qing-Song Zhou, Chao-Dong Zhu, Simon Y.W. Ho

## Abstract

Passerine birds are among the most diverse and species-rich groups of vertebrates, but the timescale of their evolution has been difficult to resolve with confidence. The fossil record of early passerines is relatively sparse and molecular-clock estimates of the passerine crown age have varied widely, with most previous studies relying on external fossil calibrations or assumptions relating to Gondwanan vicariance. In this study, we estimated the passerine evolutionary timescale by incorporating a set of 43 passerine fossils selected through a detailed assessment, while using a Bayesian tip-dating approach with the unresolved fossilized birth-death process. Our analyses ultimately place the passerine crown age in the Eocene, which largely closes the gap between molecular and palaeontological estimates of the passerine evolutionary timescale. Our date estimates are somewhat influenced by the prior probability density for the starting time of the diversification process. Through a simulation study, we show that the effect of the starting-time prior can be attenuated by the inclusion of morphological data for fossil and extant taxa. Overall, our study demonstrates that incorporating a curated, comprehensive set of fossils is effective in producing a well-resolved estimate of the passerine evolutionary timescale, while highlighting potential avenues for refining this estimate using Bayesian tip-dating analyses.

## 1. Introduction

Passerine birds form the largest avian order (Aves: Passeriformes), with over 6700 species in three suborders: Acanthisitti (New Zealand wrens), Tyranni (suboscines), and Passeri (oscines or songbirds) (Gill et al. 2024). Despite their enormous species diversity, ubiquity, and an evolutionary history spanning tens of millions of years, the early fossil record of passerines is sparse compared with those of other major bird groups (Mayr 2022). Consequently, their evolutionary timescale has largely been inferred using molecular dating. Some studies have placed the origin of crown passerines in the Cretaceous (e.g., Brown et al. 2008; Stervander et al. 2020) or around the Cretaceous–Paleogene boundary (e.g., Ericson et al. 2014; Selvatti et al. 2015), whereas phylogenomic analyses have estimated an Eocene origin (Jarvis et al. 2014; Prum et al. 2015; Oliveros et al. 2019; Stiller et al. 2024) (Fig. 1). A recent total-evidence dating analysis of combined molecular and morphological data, which included nine fossil taxa, also found an early Eocene age for crown passerines (Lowi-Merri et al. 2024). Within Passeriformes, molecular studies have inferred a middle Eocene to early Oligocene crown age for songbirds (i.e., the suborder Passeri), which are believed to have originated on the Australian continental plate (e.g., Prum et al. 2015; Moyle et al. 2016; Stiller et al. 2024). Thus, resolving the timescale of passerine evolution has implications for understanding the impacts of the Cretaceous–Paleogene mass extinction, the biogeographic effects of Gondwanan fragmentation, and the conditions that led to the successful diversification of the largest order of birds (e.g., Ericson et al. 2003; Selvatti et al. 2015; Crouch and Tobias 2022; McCullough et al. 2022).

**Figure 1.**
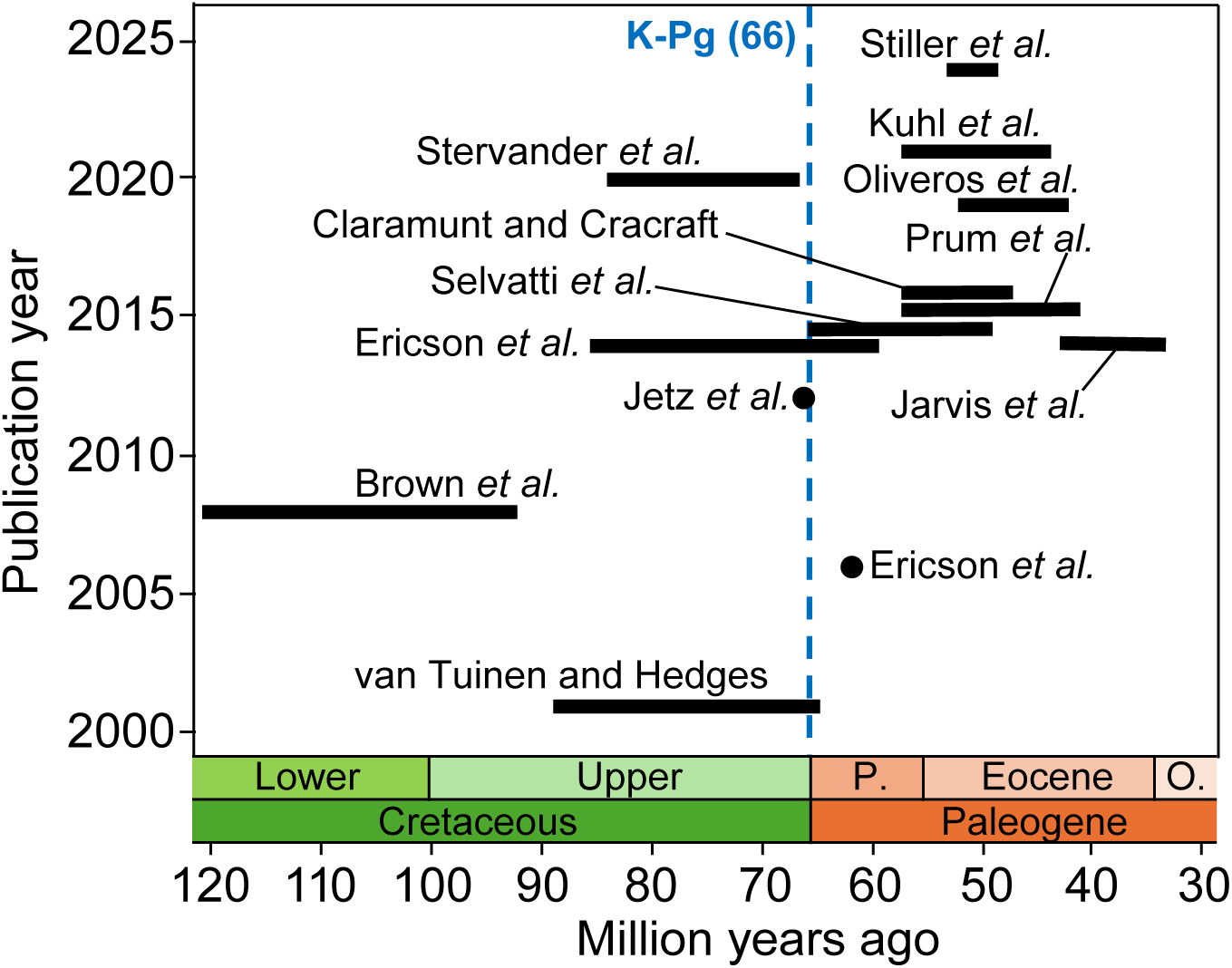
Estimates of the passerine crown age inferred in selected previous studies. Date estimates are represented by bars (95% credible intervals [CIs]) or dots (point values) along the *x*-axis, whereas publication years of the date estimates are shown on the *y*-axis. Note that we present only a single date estimate from each study, even for those that reported multiple estimates.

Molecular dating analyses of passerines have typically been calibrated using constraints or informative priors on the ages of internal nodes (Drummond et al. 2006; Yang and Rannala 2006). However, these methods are not well suited to the study of passerines, because the sparse fossil record of these birds is unable to support informative age constraints across the phylogeny. As a consequence, some molecular dating analyses of passerines have relied on secondary calibrations (based on previous molecular estimates; e.g., Jønsson et al. 2011; Gibb et al. 2015) or on the biogeographic assumption that the basal divergence in passerines, between New Zealand wrens and all other passerines, coincided with the split between New Zealand and Antarctica (e.g., Barker et al. 2004; Ericson et al. 2014; Selvatti et al. 2015). By contrast, Bayesian tip-dating approaches can provide a more effective approach to reconstructing the passerine evolutionary timescale, because these methods allow fossils to be included as sampled taxa while providing time calibrations for the molecular clock (Ronquist et al. 2012; O’Reilly et al. 2015; Gavryushkina and Zhang 2020). A significant step in the evolution of Bayesian tip dating was the development of the fossilized birth-death (FBD) process, which provides a tree prior that accounts for fossil recovery along a history of speciation and extinction (Stadler 2010; Heath et al. 2014; Zhang et al. 2016). The placement of each fossil can then be inferred from morphological data (i.e., resolved FBD trees) or left unresolved within any user-specified clade constraints (i.e., unresolved FBD trees; Heath et al. 2014).

Bayesian tip dating on resolved FBD trees can be performed using combined data sets of molecular and morphological characters, an approach known as total-evidence tip dating (Zhang et al. 2016; Gavryushkina et al. 2017). However, this method has limited application in groups with a paucity of well-preserved fossil taxa, as exemplified by the recent study of a passerine data set that comprised 47 extant and only nine fossil taxa (Lowi-Merri et al. 2024). The skeletal characters of passerines also show relative uniformity across species, as well as many instances of convergence, which present considerable difficulties for constructing informative morphological data sets from fossil taxa (Mayr 2013; Stervander et al. 2020).

These difficulties are compounded by the logistical challenges of compiling morphological data for such a species-rich clade. As an alternative, Bayesian tip dating can be performed on unresolved FBD trees, where morphological characters do not need to be coded for fossil taxa; this approach uses molecular sequence data exclusively for extant taxa, while the placements of fossil taxa are unresolved and are only subject to phylogenetic constraints that are specified by the user (Heath et al. 2014). However, both tip-dating approaches require the FBD tree prior to be conditioned on an informative starting time (Gavryushkina and Zhang 2020), which is typically either the origin time of the FBD process (*t*_o_) or the time to the most recent common ancestor (i.e., the root age; *t*_mrca_) and effectively serves as an internal node calibration. Specifying the prior for the timing of these events is typically a difficult exercise because the fossil record cannot offer direct information about these dates. Furthermore, the impact of the starting-time prior remains poorly characterized, despite previous evaluations of the performance of Bayesian tip dating under a range of simulation conditions (Luo et al. 2020, 2023). This represents a substantial gap in knowledge, given that calibration priors at internal nodes, including the root, can be highly influential on molecular estimates of evolutionary timescales (e.g., Drummond et al. 2006; Ho and Phillips 2009; Duchêne et al. 2014; Warnock et al. 2015).

In this study, we infer the evolutionary timescale of passerine birds using Bayesian tip dating on unresolved FBD trees, which is unable to handle genome-scale data sets but makes greater use of fossil information compared with the node-dating methods that have been used for previous estimates of the passerine evolutionary timescale. We assemble a curated, comprehensive set of fossil occurrences (Table 1), using a strict set of criteria to assess and select these from all of the fossil taxa referred to Passeriformes (Gandolfo et al. 2008; Parham et al. 2012). Our Bayesian tip-dating analyses are performed on a multilocus data set, with genus-level and family-level sampling of taxa. We also test the impact of the starting-time prior under a range of conditions, allowing us to evaluate the robustness of our date estimates. Our study supports an Eocene crown age for passerine birds and suggests that the date estimate can be refined by the addition of morphological data.

**Table 1.**
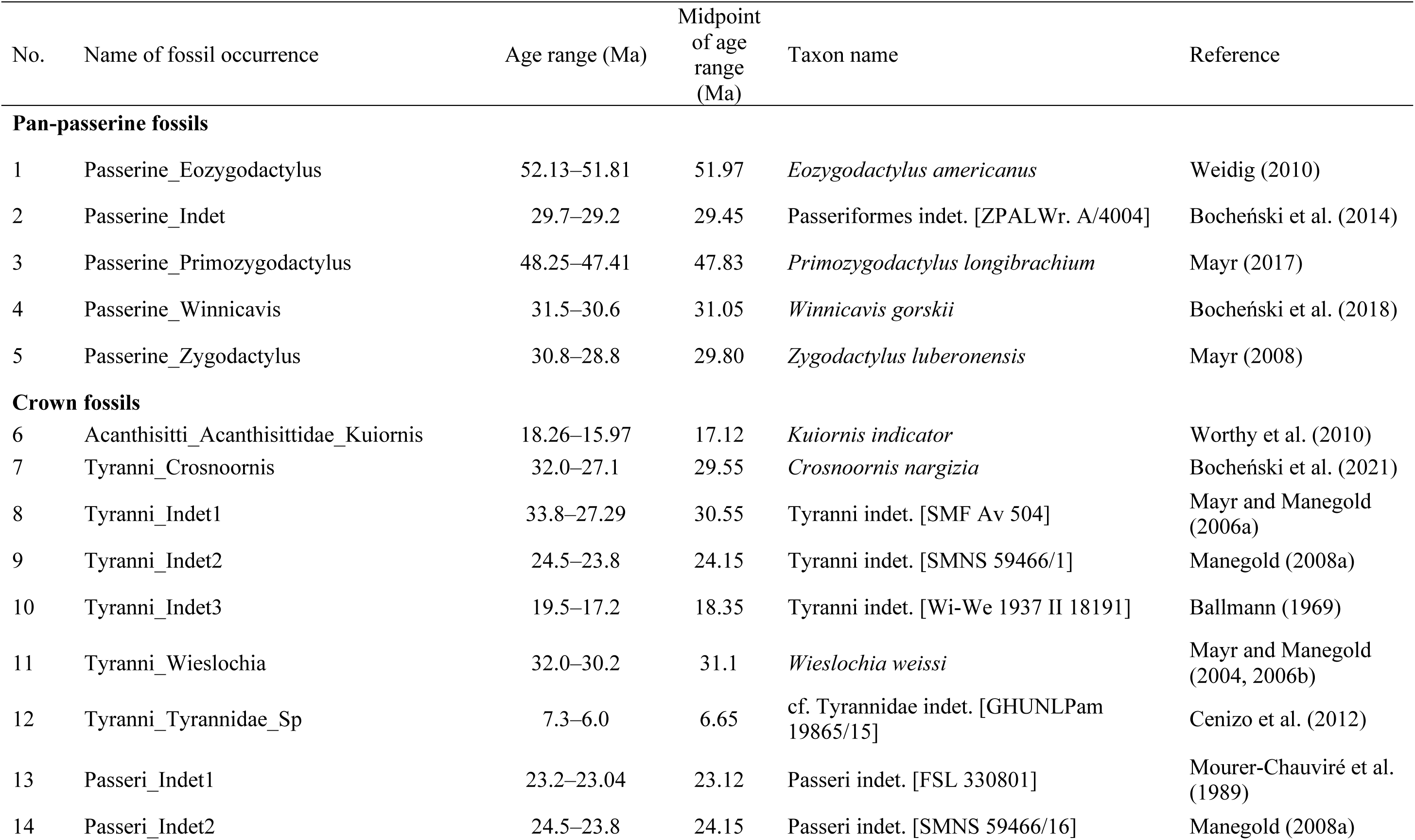
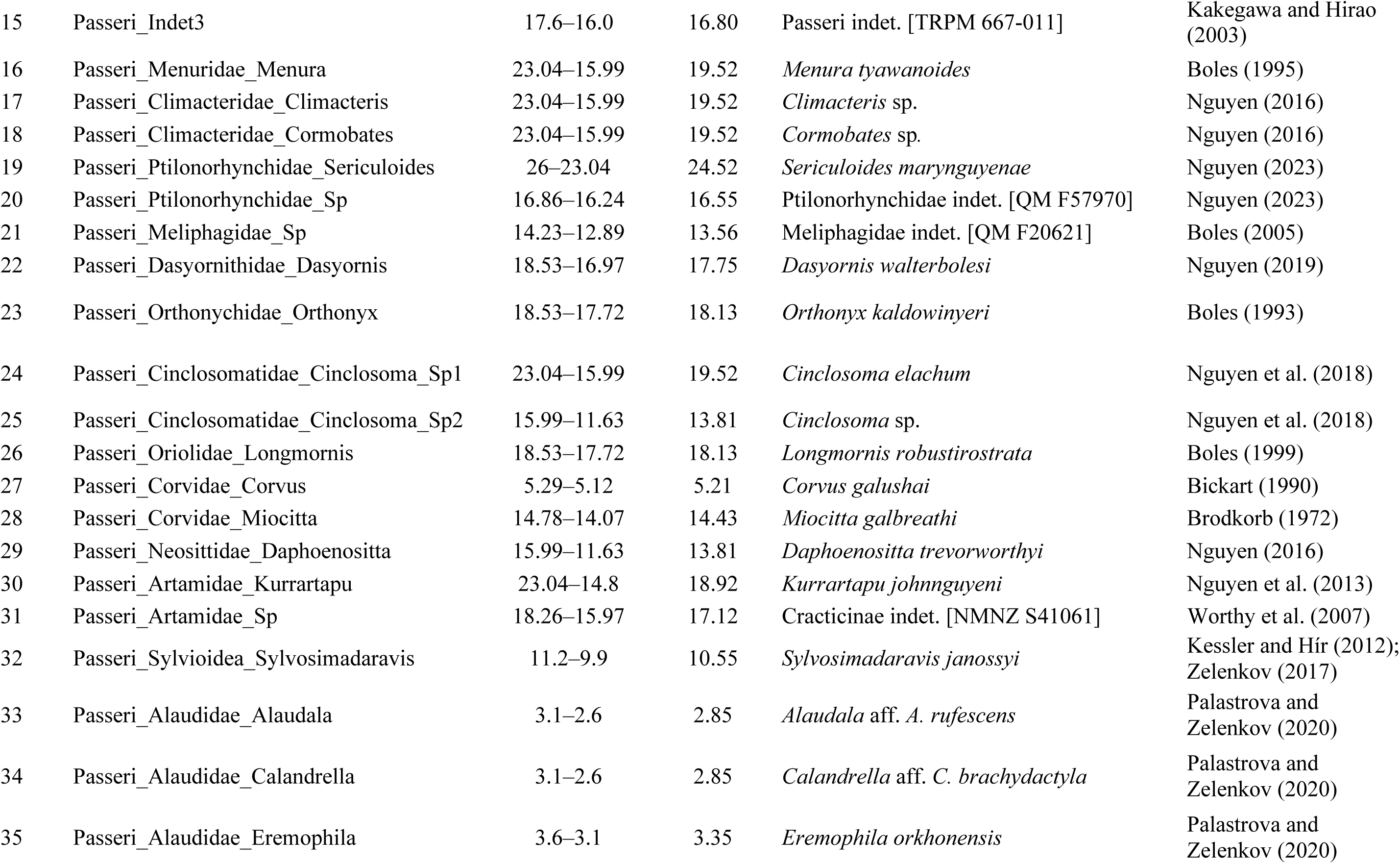
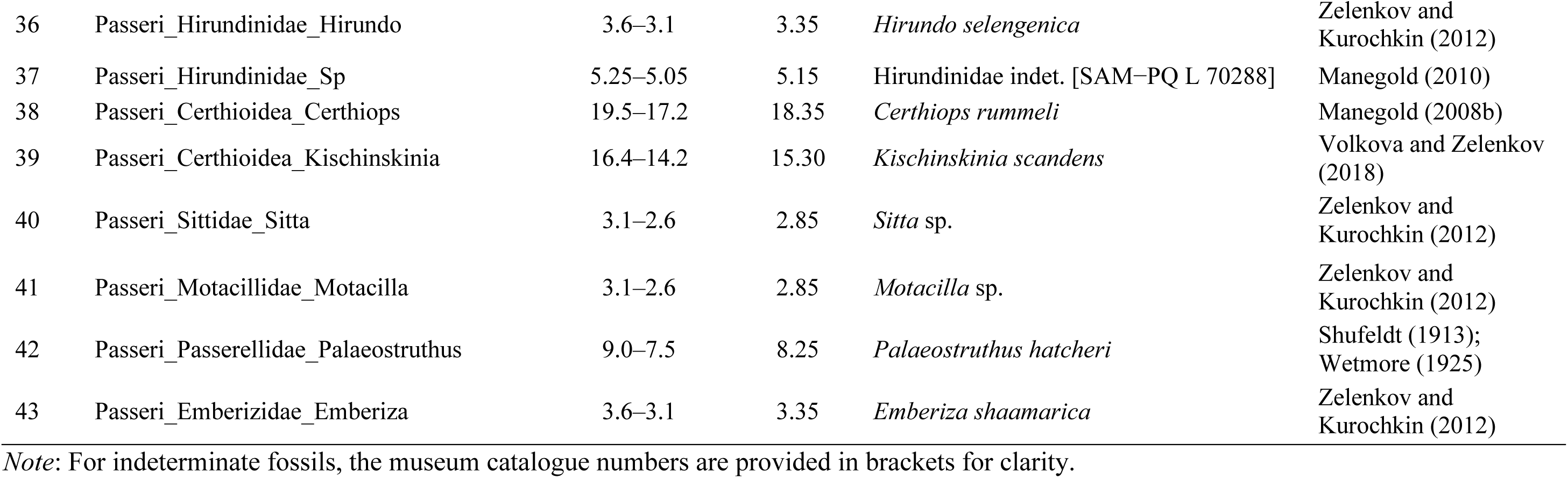
Fossil taxa used in our analyses of passerine birds.

## 2. Materials and Methods

### 2.1. Molecular Data and Fossil Occurrences

We adopted a multilocus data set that was assembled in a previous phylogenetic study of passerine birds (Selvatti et al. 2015). This data set was chosen for a combination of taxonomic representation and computational tractability, and comprises the nucleotide sequences of five mitochondrial genes (*CYTB*, *COX1*, *ND2*, *12S*, and *16S*), two nuclear genes (*RAG1* and *RAG2*), and two introns (*MYO* and *ODC*). Following Selvatti et al. (2015), we used the pruned data set from a total of 1119 extant genera, but we included only 322 ingroup genera (from an initial set of 329 genera) after excluding taxa with large proportions of missing data. We also filtered the sequences to produce a concatenated alignment of 9704 nucleotides. This formed our genus-level data set, whereby each taxon represented one extant genus of passerine birds (for some taxa, the sequence data are from a mosaic of congeneric species). We then obtained a family-level data set by selecting 138 taxa that each represented one extant family of passerine birds. We updated the taxonomic information of a few taxa in accordance with recent work (Gill et al. 2024), and our analyses were performed using both the genus- and family-level data sets.

In our data analyses, we considered all of the fossil taxa that have been referred to Passeriformes. After evaluating candidate fossils using a strict set of criteria (Gandolfo et al. 2008; Parham et al. 2012), we chose to include a total of 43 fossil occurrences, including 31 fossil taxa from Passeri, six from Tyranni, and one from Acanthisitti (Table 1; Fig. 2a). Most of these taxa have been identified to at least family-level (some even to species, such as *Orthonyx kaldowinyeri*; Nguyen et al. 2014), leaving six taxa (three each from Passeri and Tyranni) without further taxonomic resolution *a priori*. In addition to these crown-group fossils, we included five passerine fossils that were unable to be constrained to the crown group, which included three stem passerines (Mayr 2008, 2017; Weidig 2010) and two fossils with unresolved affinities (Bocheński et al. 2014, 2018) (Table 1). Henceforth, we refer to these five as “pan-passerine” fossils.

**Figure 2.**
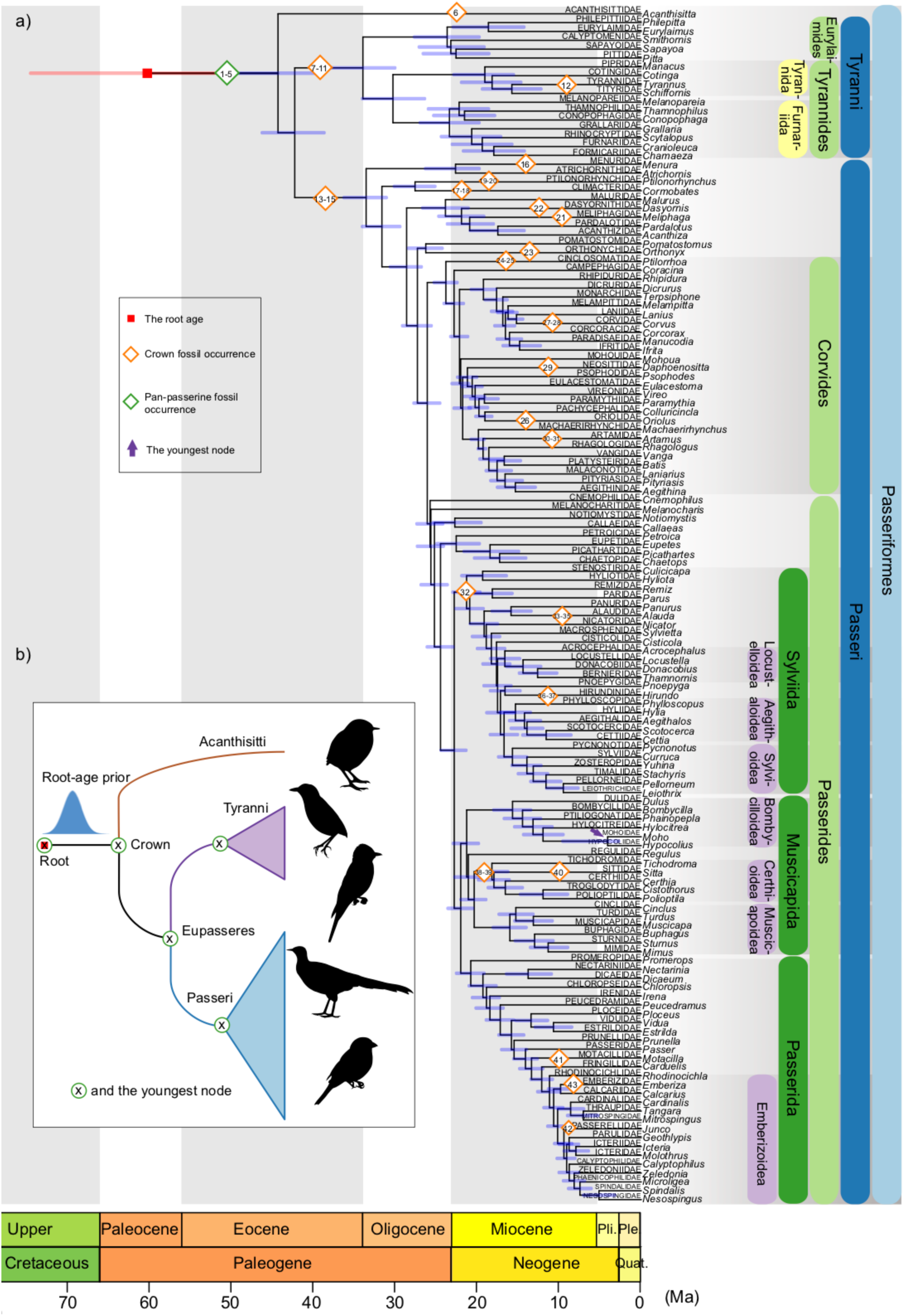
a) Maximum-clade-credibility tree of passerine birds from our analyses with monophyly constraints for extant taxa at the family level under the uniform(51.97,90.9) root-age *t*_mrca_ prior. Tree topology was informed by phylogenomic analyses (Oliveros et al. 2019; Stiller et al. 2024), whereas posterior medians and 95% CIs (blue horizontal bars) of node times were estimated by our analyses. Fossil occurrences are denoted across the tree according to their taxonomic placements and assumed monophyly of relevant passerine groups, with their numbers corresponding to those in Table 1 and their placements along the branches being arbitrary (not reflecting specific time information). Due to monophyly constraints for extant taxa, note that the taxonomic placement of fossil 42 here results in inconsistency of its topological constraint between our analyses with and without monophyly constraints for extant taxa. b) A schematic showing the key time points (denoted by “X” in green circles) that we focused on when we evaluated the effects of the choice of the *t*_mrca_ prior on Bayesian tip-dating analyses. Bird silhouette images are from Phylopic (phylopic.org) and are in the public domain.

We used the taxonomic descriptions of the fossils to set clade constraints in our phylogenetic analyses. Although some of the fossil taxa have a degree of age uncertainty, we simply adopted the midpoints of their age ranges, which had median ages ranging from 2.85 Ma (e.g., Palastrova and Zelenkov 2020) to 51.97 Ma (*Eozygodactylus americanus*; Weidig 2010). Justifications for these fossil calibrations are presented in a format adapted from that of Ramírez-Barahona et al. (2020).

### 2.2. Bayesian Tip-dating Analyses

We performed Bayesian molecular tip-dating analyses using the nucleotide sequences and fossil ages described above. Given that the sequence data were sampled with the aim of including representatives at the genus or family level, we used the FBD process accounting for diversified sampling of extant species as the tree prior (Zhang et al. 2016). For the FBD model parameters, which are a reparameterization of constant rates of speciation (*λ*), extinction (*μ*), and fossil recovery (*ψ*) (Heath et al. 2014), we assigned an exponential(1.0) prior for diversification rate *d* = *λ* – *μ*, beta(0,1) prior for turnover *r* = *μ* / *λ*, and beta(1,10) prior for fossil sampling proportion *s = ψ* / (*ψ* + *μ*). We applied a topological constraint for each fossil based on its taxonomic placement and assumed monophyly of the corresponding passerine group; for the sake of comparison, we ran additional analyses with the five pan-passerine fossils excluded.

The molecular data were partitioned into five subsets: (i) first and second codon sites of mitochondrial *COX1*, *CYTB*, and *ND2*; (ii) first and second codon sites of nuclear *RAG1* and *RAG2*; (iii) third codon sites of nuclear *RAG1* and *RAG2*; (iv) mitochondrial *12S* and *16S* rRNA; and (v) *MYO* and *ODC* introns. Each subset was assigned a separate GTR+G model.

A single uncorrelated lognormal relaxed-clock model was applied to the molecular data, with a uniform(0,1) prior for the mean evolutionary rate. For analyses at the genus and family levels, we fixed the extant species’ sampling fraction *ρ* at 0.0496 and 0.0213, respectively, assuming a total of ∼6500 extant passerine species (Gill et al. 2024).

The FBD tree prior needs to be conditioned on a starting time, so we first assigned a uniform prior for the root age *t*_mrca_. The minimum bound of this uniform prior matched the age of the oldest fossil: the 51.97-Myr-old *Eozygodactylus americanus* (see Weidig 2010) when the five pan-passerine fossils were included; or the 31.1-Myr-old *Wieslochia weissi* (see Mayr and Manegold 2004, 2006a) when the pan-passerine fossils were excluded. The maximum bound was set to 90.9 Ma: chosen as a random time point older than the maximum age of the New Zealand–Antarctica split 85 Ma, but younger than the Cretaceous–Jurassic boundary 145 Ma (McLoughlin 2001; Zachos et al. 2001). Second, we applied a uniform *t*_mrca_ prior with a maximum bound of 85 Ma and a minimum bound of 52 Ma; this corresponds to the time range of the New Zealand–Antarctica split, which is believed to have had an impact on early passerine diversification (McLoughlin 2001; Zachos et al. 2001). Third, we applied a uniform(51.97,145) or uniform(31.1,145) prior on *t*_mrca_, with the maximum bound being the boundary between the Cretaceous and Jurassic. Finally, we specified normal(53.5,10) and normal(83.5,10) priors on *t*_mrca_, corresponding to different hypotheses regarding the timing of the New Zealand–Antarctica vicariance (Ericson et al. 2014) (Table 2).

**Table 2.**
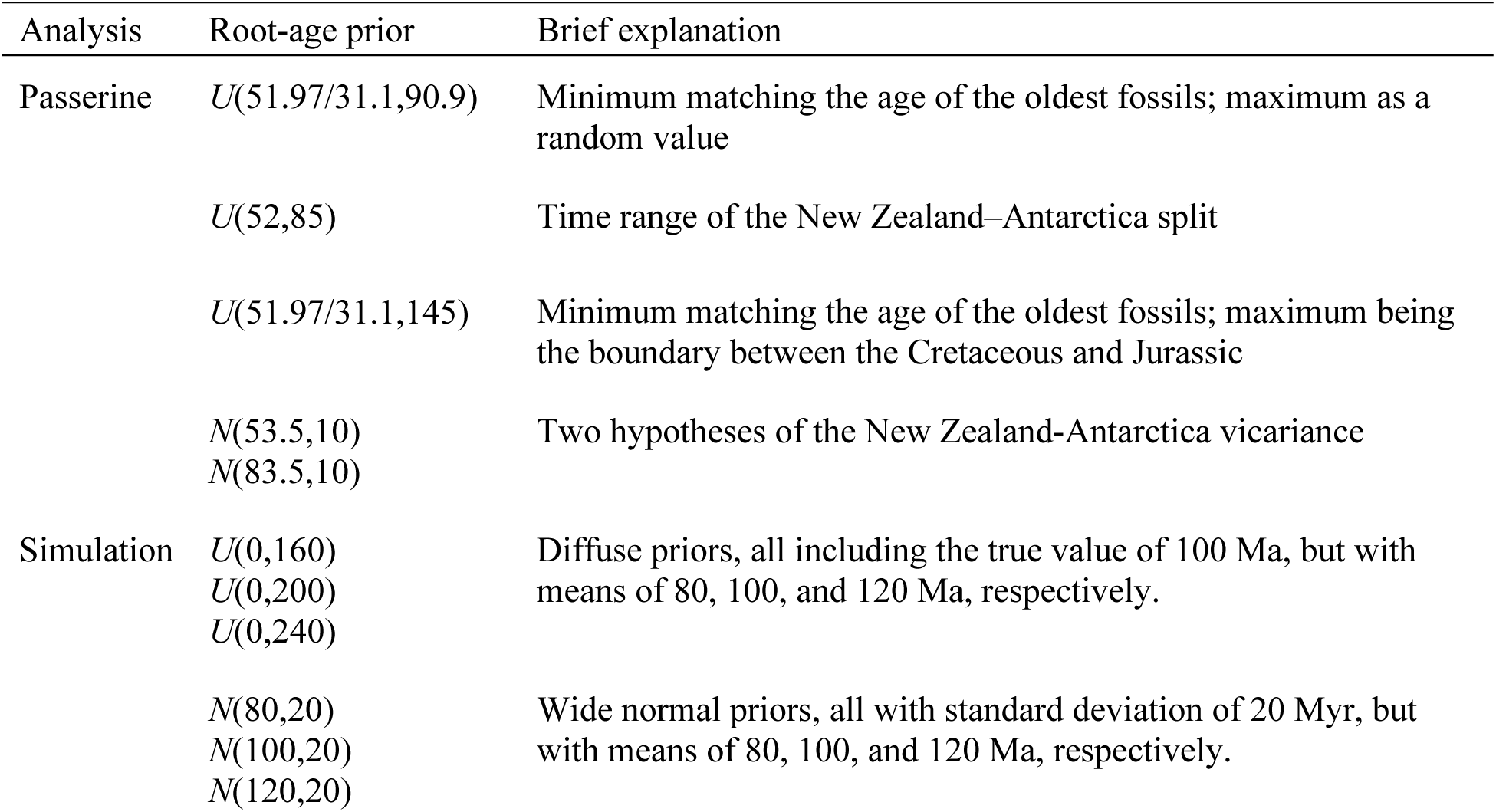
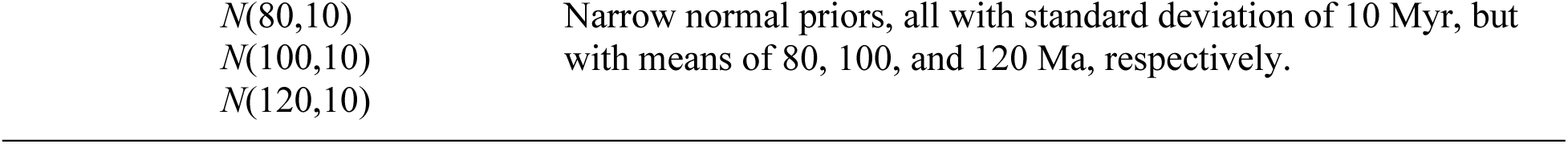
Settings of the root-age prior in our passerine and simulation analyses. Analysis Root-age prior Brief explanation.

Our tip-dating analyses on unresolved FBD trees were performed using BEAST v2.6 (Bouckaert et al. 2019) with the BDSKY v.1.4.5 package (Stadler et al. 2013). All posterior distributions were estimated by Markov chain Monte Carlo (MCMC) sampling. For each of the 20 analyses (2 taxonomic levels × 5 root-age priors × 2 fossil sets), we carried out four independent MCMC runs, with samples drawn every 10,000 steps over a total of 100 or 200 million steps. We adjusted the burn-in fractions and checked for convergence across chains using Tracer v1.7 (Rambaut et al. 2018). Sufficient sampling was confirmed using LogAnalyzer v2.6. To enforce the position of the root, in addition to the topological constraints on the fossils, we placed a monophyly constraint on Eupasseres (Passeri + Tyranni). For each MCMC analysis, we identified the maximum-clade-credibility tree among the sampled trees (at least 22,000 trees) using TreeAnnotator v2.6, with fossil taxa pruned beforehand using the application FullToExtantTreeConverter.

With topological constraints for fossils and a monophyly constraint on Eupasseres, the above analyses jointly estimated node times and tree topologies. As a comparison, we repeated all of the analyses with not only topological constraints on the fossils, but also monophyly constraints for 135 extant taxa at the family level (i.e., three fewer taxa than our original data, due to uncertainty in their phylogenetic position) based on the relationships inferred in two phylogenomic analyses (Oliveros et al. 2019; Stiller et al. 2024). The topology of extant taxa was then fixed for the estimation of node times (Fig. 2a). All other settings for these 10 tip-dating analyses (5 root-age priors × 2 fossil sets) were as described above.

### 2.3. Simulation analyses

Synthetic data were obtained primarily from a previous study (Luo et al. 2023). In brief, 100 complete species trees were generated with crown ages *t*_c_ equal to the root age *t*_mrca_ of 100 Ma. Along these trees, 2–15 (median 7) fossil occurrences were sampled under a Poisson process with a recovery rate *ψ* = 0.003. Rate variation among lineages was simulated using an uncorrelated lognormal relaxed clock (mean 10^-3^ substitutions/site/Myr, standard deviation 2×10^-4^) (Drummond et al. 2006), with the nucleotide sequences (five ‘loci’, each with a length of 1000 bp) of extant taxa evolving under an HKY+G substitution model. In contrast with the approach taken by Luo et al. (2023), we adopted both diversified and random sampling strategies for extant taxa (Höhna et al. 2011), with sampling fraction *ρ* = 0.1, 0.5, and 1.0. For dating analyses under diversified sampling, nucleotide sequences of the *n* extant taxa were combined with the fossil occurrences with dates at least as old as the (*n* − 1)th oldest divergence in the reconstructed tree; for those under random sampling, nucleotide sequences were analysed along with all sampled fossil occurrences.

In addition to the data above, we included morphological characters for both extant taxa and sampled fossils for total-evidence tip dating. Rate variation among lineages was simulated using the same settings as for the molecular data. Morphological characters evolved under the Mkv model (Lewis 2001), primarily producing data sets comprising 500 four-state characters. After sampling fossil taxa according to sampling fraction *ρ* and (where appropriate) the (*n* − 1)th oldest divergence, we removed any invariant characters.

For either total-evidence tip dating (with morphological characters and nucleotide sequences) or tip dating on unresolved FBD trees (with nucleotide sequences only), we generally matched models to those used for simulation, including the FBD model accounting for diversified or random sampling. Our prior settings followed those of Luo et al. (2023), including diffuse priors for the FBD model parameters, true values for sampling fraction *ρ*, true ages of fossil occurrences, separate relaxed-clock models for the molecular and morphological data for total-evidence tip dating, and a single clock model for tip dating on unresolved FBD trees. For tip dating on unresolved FBD trees, we placed a topological constraint on each fossil based on its parent node in the true FBD trees.

For the FBD starting time, we explored variations of the root-age *t*_mrca_ prior for both tip-dating approaches. First, we adopted diffuse uniform(0,160), uniform(0,200), and uniform(0,240) priors for *t*_mrca_, all of which include the true value 100 Ma, but have respective means of 80 Ma, 100 Ma, and 120 Ma. Second, we used wide normal priors, with means of 80 Ma, 100 Ma, and 120 Ma, with standard deviations of 20 Myr (i.e., 20% of the true *t*_mrca_). Third, we used narrow, more informative normal priors with means of 80 Ma, 100 Ma, and 120 Ma, but with standard deviations of 10 Myr. Thus, our simulation study involved a total of 10,800 analyses (100 species trees x 3 extant sampling fractions x 2 sampling strategies x 2 tip-dating approaches x 9 *t*_mrca_ priors).

Dating analyses were similar to those described above for the passerine data and followed Luo et al. (2023). To evaluate the date estimates, we focused on posterior estimates of the root age *t*_mrca_, the crown age *t*_c_, and the tree length (i.e., sum of branch lengths). We also examined the posterior estimates of total evolutionary change (the product of clock rate and tree length) and clock rate, as well as the Robinson-Foulds topological distance between the maximum-clade-credibility trees and the true topologies (Robinson and Foulds 1981).

## 3. Results

### 3.1. Evolutionary History of Passerine Birds

Our Bayesian tip-dating analyses on unresolved FBD trees collectively placed the origin of crown Passeriformes in the early-middle Eocene, with posterior medians of ∼43–53 Ma (95% credible intervals [CIs] of ∼39–57 Ma) (Fig. 2a and 3). We dated the divergence between Passeri and Tyranni to the middle Eocene, with posterior medians of ∼41–48 Ma (95% CIs of ∼38–53 Ma), and with these two crown groups emerging in the late Eocene (posterior medians ∼33–38 Ma and ∼34–39 Ma, respectively). Our analyses generally supported Paleogene emergences for the major passerine clades, with infraorders such as Tyrannides, Corvides, and Passerides being dated to the Oligocene (Fig. 2a), whereas the divergences between families and genera generally occurred in the Neogene with posterior medians of >2.97 Ma for the youngest node (e.g., the divergence between the families Mohoidae and Hypocoliidae in Fig. 2a). Additionally, posterior medians of the root age (i.e., the time to the most recent common ancestor of passerine birds) ranged from 43 to 69 Ma (95% CIs of ∼39–86 Ma), spanning from the middle Eocene to late Cretaceous (Fig. 2a and 3).

**Figure 3.**
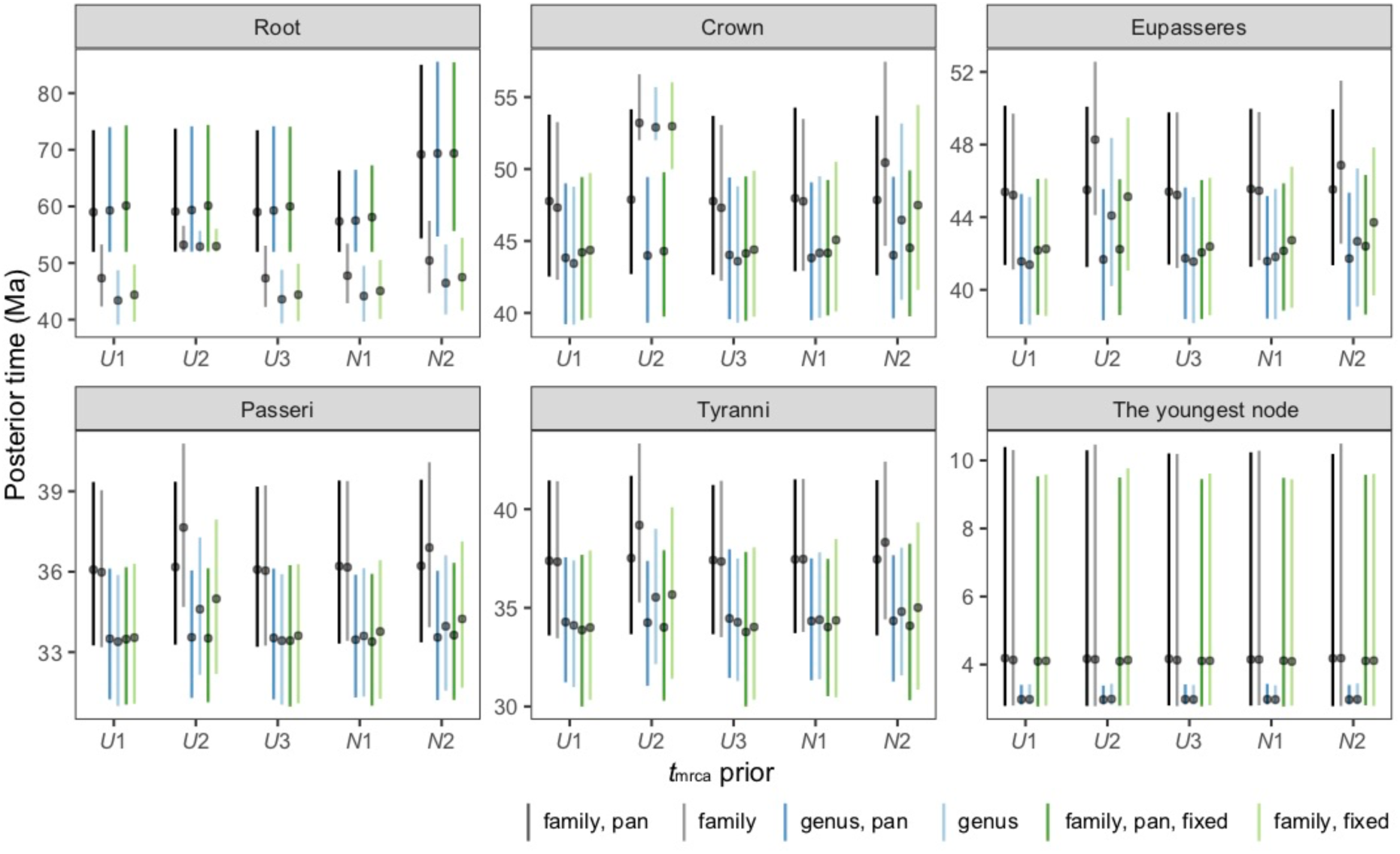
Posterior time estimates from our analyses of passerine birds. For each key time point, posterior medians (black circles) and 95% CIs (lines) are shown for analyses at either family (grey lines) or genus level (blue lines) under each root-age *t*_mrca_ prior (*x*-axis), with pan-passerine fossils included (dark coloured lines) (denoted by “pan” in the legend) or excluded (light coloured lines); additional estimates from analyses with hierarchical monophyly constraints for extant taxa at the family level are denoted by green lines (denoted by “fixed” in the legend). For the sake of clarity, we use *U*1, *U*2, *U*3, *N*1 and *N*2 to represent the uniform(51.97/31.1,90.9), uniform(52,85), uniform(51.97/31.1,145), normal(53.5,10), and normal(83.5,10) *t*_mrca_ priors, respectively.

Our date estimates for the passerine evolutionary timescale were obtained under several different priors for the root age *t*_mrca_, variously informed by the age of the oldest fossil included in the analysis, hypotheses of the timing of the New Zealand–Antarctica vicariance event, and other sources (Table 2). By focusing on several representative nodes in the phylogeny, we also evaluated the robustness of our Bayesian tip-dating estimates to the inclusion or exclusion of five pan-passerine fossil occurrences, the level of taxon sampling, and the inclusion or omission of monophyly constraints for extant taxa (Fig. 3). For analyses without monophyly constraints for extant taxa, when the five pan-passerine fossils were included, varying the *t*_mrca_ prior only influenced estimates of the root age; the uniform *t*_mrca_ priors led to moderate estimates of the root age, whereas the normal priors resulted in estimates that recapitulated the prior expectations (reflected by the mean values). When the pan-passerine fossils were excluded, root-age *t*_mrca_ priors influenced the date estimates of all focal nodes except the youngest node; the uniform(52,85) and normal(83.5,10) *t*_mrca_ priors led to larger posterior medians. We found that including or excluding the pan-passerine fossils primarily affected estimates of the root age (with a maximum difference of 23 Myr); our analyses at the genus level generally yielded younger date estimates (< 4.2 Myr) with narrower 95% credible intervals than their counterparts at the family level, except for similar estimates of the root age in analyses with the pan-passerine fossils included (Fig. 3).

For analyses with monophyly constraints for extant taxa, the effects of the choice of *t*_mrca_ prior displayed patterns similar to those described above (Fig. 3). In particular, including the pan-passerine fossils led to older estimates of the root age, but somewhat younger or similar estimates of other node ages, than when these fossils were excluded. Compared with their counterparts (i.e., analyses without monophyly constraints for extant taxa), most date estimates were younger, but patterns for estimates of deep nodes (e.g., the crown age) varied with specific *t*_mrca_ priors and the inclusion or exclusion of pan-passerine fossil occurrences.

We found a maximum gap of 3.8 Myr between estimates from these analyses and their counterparts.

### 3.2. Evaluation Using Synthetic Data

In our simulation analyses, we found that the posterior estimates of the root age *t*_mrca_ were very close to those of the crown age *t*_c_, so we focus on *t*_c_ hereafter. The root-age *t*_mrca_ prior primarily affected the posterior estimates of *t*_c_ and tree length, among the various estimates that we monitored.

In the analyses on unresolved FBD trees, we found that the choice of *t*_mrca_ prior had an effect on the estimates of *t*_c_ (Fig. 4a). With uniform *t*_mrca_ priors, *t*_c_ tended to be overestimated. With normal priors, posterior medians of *t*_c_ were generally close to the prior expectations, but the wide normal priors led to estimates that were older and more diffuse than those obtained using the narrower priors (e.g., normal(80,20) versus nomal(80,10)). Estimates of *t*_c_ under the normal priors were more precise than those under the uniform priors. Estimates of the tree length exhibited similar patterns to those of *t*_c_. Compared with the *t*_mrca_ prior, taxon-sampling density (as with the genus- and family-level sampling in the analyses of the passerine data) showed limited impacts on date estimates, with the largest effects seen in analyses under uniform priors (e.g., an increase in date estimates). Date estimates from analyses with diversified taxon sampling showed no difference in patterns of accuracy from those with random sampling. However, the latter analyses showed precision improving with denser taxon sampling, in contrast with that seen in the former analyses.

**Figure 4.**
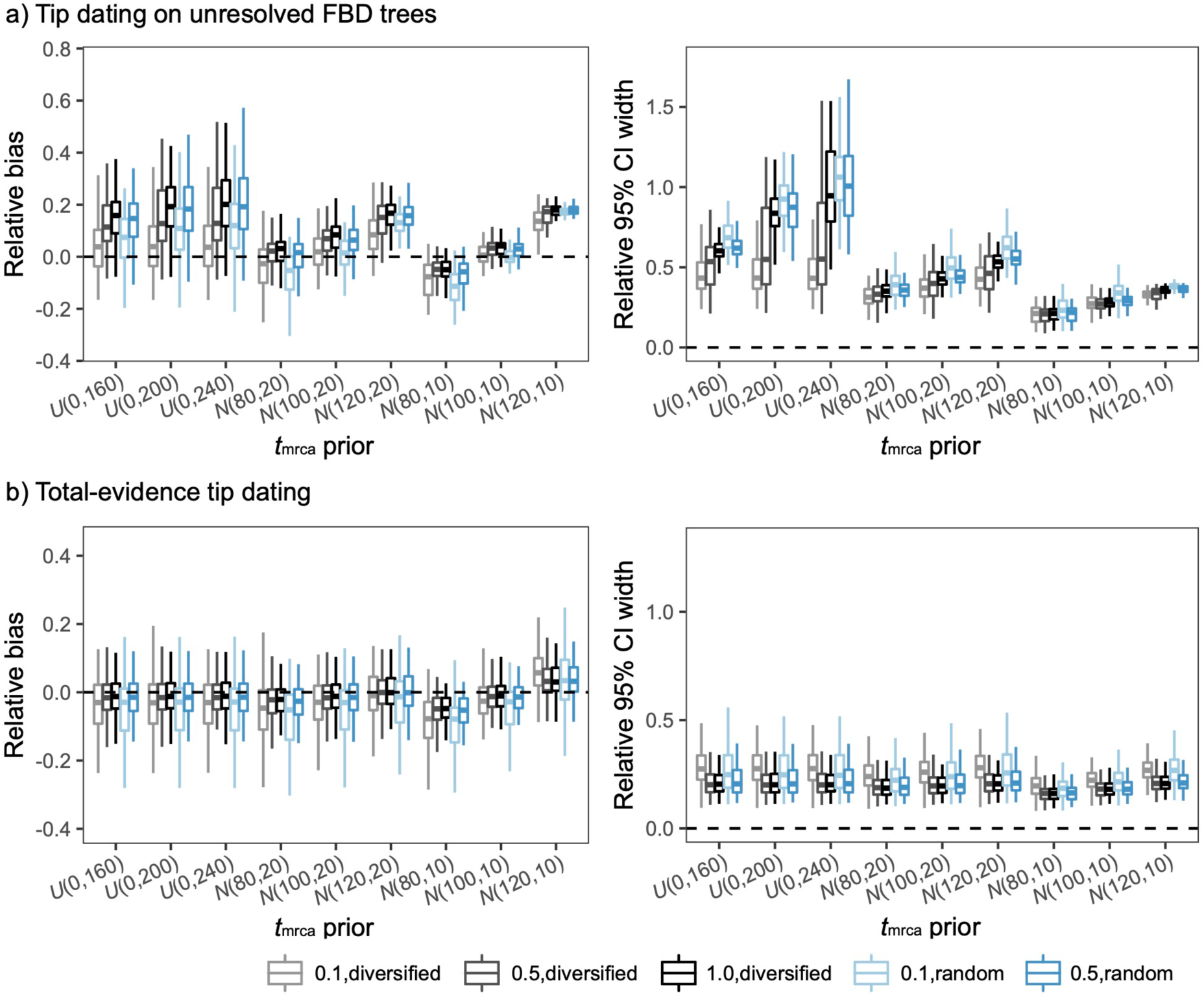
Posterior estimates from the simulation analyses. a) Boxplots summarizing estimates of the crown age *t*_c_ from tip dating on unresolved FBD trees, with accuracy measured by relative bias (distance between the posterior median and the true value, divided by the true value) and precision by relative 95% CI width (posterior 95% CI width divided by the true value). b) Boxplots summarizing estimates of *t*_c_ from total-evidence tip dating. For both (a) and (b), 100 replicates are summarized into an independent boxplot at each level (extant taxon sampling fraction *ρ* = 0.1, 0.5, and 1.0 from left to right, in increasingly dark coloured lines) of diversified (black) or random (blue) taxon sampling under each root-age *t*_mrca_ prior (*x*-axis). Dashed horizontal lines indicate the true or expected values. Note that when *ρ* = 1, random sampling does not apply, as random sampling is then identical to diversified sampling.

Compared with the analyses on unresolved FBD trees, the analyses that included morphological data showed that posterior estimates of *t*_c_ were generally accurate and robust to the choice of the *t*_mrca_ prior; there was only a slight dependence on the informativeness of the specific prior (Fig. 4b). Increasing taxon-sampling density had smaller impacts than those seen in analyses on unresolved FBD trees, but with greater precision in date estimates. There was no evident disparity between date estimates from analyses with random and diversified taxon sampling.

## 4. Discussion

In this study, we have presented a Bayesian tip-dating estimate of the passerine evolutionary timescale that incorporates time information from an extensive set of fossil occurrences. The 43 fossils represent the largest set of fossil taxa that has been incorporated into a molecular dating analysis of passerine birds. The results of our analyses firmly place the origin of crown Passeriformes in the Eocene (43–53 Ma). Thus, our inferred timescale is consistent with some recent estimates using fossil calibrations (Oliveros et al. 2019; Stiller et al. 2024) and total-evidence dating (Lowi-Merri et al. 2024), but conflicts with other estimates calibrated by fossils (Brown et al. 2008; Stervander et al. 2020) or by the New Zealand–Antarctica split (Ericson et al. 2014; Selvatti et al. 2015). The emerging consensus of an Eocene origin for crown passerines lays the foundation for interpreting the evolutionary history of the morphological and ecological traits that contributed to their successful diversification (Crouch and Tobias 2022; Miles et al. 2023). It also provides a dated phylogenetic framework for detailed biogeographic analyses of passerine birds (e.g., Ericson et al. 2014; Oliveros et al. 2019) and the drivers of their evolutionary rates (Duchêne et al. 2025).

Our estimates suggest that, following the split between Acanthisitti and Eupasseres, some of the major divergences among passerine clades occurred in the Paleogene and particularly within a small time window around the Eocene and Oligocene, indicating relatively rapid diversification of these clades. The timescale inferred here also suggests that shifts in diversification rate have occurred across clades of passerines at finer scales, which have ultimately led to marked disparities in species richness across major clades (Ricklefs 2003), such as a much larger number of species within Passeri than within Tyranni (Gill et al. 2024). Moreover, although we have focused on the divergences among major passerine groups as representative nodes (Fig. 2b), our analyses present revised age estimates for major clades such as the infraorder Corvides, providing a framework for further research into the evolution (Jønsson et al. 2016), biogeography (McCullough et al. 2022), and morphology of these taxa (Kennedy et al. 2016).

In our Bayesian tip-dating analyses of passerine birds, the increase in fossil information has come at the cost of using a smaller molecular data set than the genome-scale data that can otherwise be analysed using node-dating methods (e.g., Jarvis et al. 2014; Oliveros et al. 2019; Stiller et al. 2024). However, this can be a reasonable trade-off under some circumstances, given that the accuracy and precision of molecular date estimates are driven primarily by the quality of the calibrating information (dos Reis and Yang 2013). Our use of a smaller molecular data set in this study has the disadvantage of reduced phylogenetic support, as shown by our analyses without monophyly constraints for extant taxa. However, this was mitigated by applying topological constraints based on phylogenomic evidence (Oliveros et al. 2019; Stiller et al. 2024). More importantly, our date estimates were reasonably robust to the inclusion or omission of monophyly constraints for extant taxa (maximum gap of 3.8 Myr), a result that underscores the greater impact of increasing the amount of fossil information.

Our analyses of passerine birds have also shown the impacts of the choice of the root-age (*t*_mrca_) prior and other factors. Compared with the limited effects of taxon-sampling density, the choice of the *t*_mrca_ prior appeared to have an influence on specific date estimates (cf. the conclusions of a recent analysis of angiosperms; Ma et al. 2025). However, these impacts depended on whether the five pan-passerine fossils were included or not. When these five fossils were included, the choice of the *t*_mrca_ prior had a noticeable effect only on estimates of the root age; but when these fossils were excluded, the choice of *t*_mrca_ prior had effects on date estimates across the tree. We attribute this difference largely to our implementation of monophyly constraints, which allows the pan-passerine fossils to jump between placements on the stem lineage or into the crown group and could restrict the influence of the starting-time prior on the date estimates of other nodes. Taken together, our results reveal that including a larger number of fossil occurrences, particularly stem fossils, can reduce the impacts of the starting-time prior on Bayesian tip dating on unresolved FBD trees (Luo et al. 2023).

As with our analyses of passerine birds with the five pan-passerine fossils excluded, our analyses using synthetic data (without stem fossil occurrences) have shown that estimates of both the crown age *t*_c_ and tree length under the unresolved FBD process are generally robust to factors such as taxon sampling and sampling strategy, but sensitive to the choice of the root-age *t*_mrca_ prior. Our analyses of synthetic and empirical data generally indicated that using a uniform prior led to stable (but not necessarily accurate) time estimates, whereas a normal prior tended to have a stronger influence on the posterior time estimates. Moreover, our simulation study reveals that the robustness of Bayesian tip dating can be improved by including morphological characters. The inclusion of morphological data allows fossil taxa to be placed in the tree more precisely, while also informing the estimates of evolutionary parameters (Luo et al. 2020). However, constructing and analysing morphological data sets can be a difficult undertaking (Lewis 2001; Sansom et al. 2010; Ronquist et al. 2016; Goloboff et al. 2019), particularly when the organisms of interest have a poor record of fossil preservation and when models of morphological character change do not fully capture the complexity of the evolutionary process (Lee and Palci 2015). Some of these challenges are heightened in studies of passerines, which have a sparse fossil record because of various taphonomic biases and display morphological uniformity and many instances of evolutionary convergence (Mayr 2013; Stervander et al. 2020). However, there have been ongoing efforts to identify diagnostic morphological characters and to resolve the taxonomic affinities of the passerine fossils that are available (e.g., Nguyen et al. 2016; Steell et al. 2023; Lowi-Merri et al. 2024).

## 5. Conclusions

We have presented an estimate of the passerine evolutionary timescale that incorporates time information from an extensive set of fossil occurrences. While firmly dating the origin of crown Passeriformes in the Eocene, our date estimates showed some sensitivity to the choice of the starting-time prior for the FBD tree prior, particularly when deep fossil occurrences were not included. This behaviour was confirmed by our further analyses of synthetic data, which also found that the addition of morphological data could improve the robustness of the date estimates. In general, our study highlights the benefits of using a curated, comprehensive set of fossil records for estimating the passerine evolutionary timescale, while raising the possibility of improving this estimate by adding morphological characters to enable a total-evidence tip-dating analysis. Along with further improvements in phylogenetic dating methods and computational capacity, we expect to see continued refinement in reconstructions of the evolutionary timescale of passerines, leading to stronger foundations for studying the diversification and biogeographic history of the largest group of living birds.

## Acknowledgements

We thank Alexandre Selvatti for providing curated nucleotide sequence data, and Hervé Sauquet for advice on justifications for fossil calibrations. TianHe-1 (A) at National SuperComputer Center in Tianjin, China, provided computing resources that have contributed to the simulation analyses performed in this paper. This work was supported by the National Science Fund for Excellent Young Scholars (32122016) and the National Natural Science Foundation of China (32470473, 32070465). We also acknowledge funding from the Australian Research Council (DE200101222 to J.M.T.N. and DP220103265 to S.Y.W.H.).

## References

1. Ballmann P. 1969. Die Vögel aus der altburdigalen Spaltenfüllung von Wintershof (West) bei Eichstätt in Bayern. Zitteliana. 1:5–60.

2. Barker F.K., Cibois A., Schikler P., Feinstein J., Cracraft J. 2004. Phylogeny and diversification of the largest avian radiation. Proc. Natl. Acad. Sci. USA 101:11040– 11045.

3. Bickart K.J. 1990. The birds of the late Miocene-early Pliocene Big Sandy Formation, Mohave County, Arizona. Ornithol. Monogr. 44:1–72.

4. Bocheński Z.M., Tomek T., Bujoczek M., Salwa G. 2021. A new passeriform (Aves: Passeriformes) from the early Oligocene of Poland sheds light on the beginnings of Suboscines. J. Ornithol. 162:593–604.

5. Bocheński Z.M., Tomek T., Świdnicka E. 2014. The first complete leg of a passerine bird from the early Oligocene of Poland. Acta Palaeontol. Pol. 59:281–285.

6. Bocheński Z.M., Tomek T., Wertz K., Happ J., Bujoczek M., Świdnicka E. 2018. Articulated avian remains from the early Oligocene of Poland adds to our understanding of Passerine evolution. Palaeontol. Electron. 21.2.32A:1–12.

7. Boles W.E. 1993. A logrunner Orthonyx (Passeriformes: Orthonychidae) from the Miocene of Riversleigh, north-western Queensland. Emu. 93:44–49.

8. Boles W.E. 1995. A preliminary analysis of the Passeriformes from Riversleigh, northwestern Queensland, Australia, with the description of a new species of lyrebird. Cour. Forschungsinst. Senckenberg.

9. Boles W.E. 1999. A new songbird (Aves: Passeriformes: Oriolidae) from the Miocene of Riversleigh, northwestern Queensland, Australia. Alcheringa. 23:51–56.

10. Boles W.E. 2005. Fossil honeyeaters (Meliphagidae) from the late Tertiary of Riversleigh, north-western Queensland. Emu. 105:21–26.

11. Bouckaert R., Vaughan T.G., Barido-Sottani J., Duchêne S., Fourment M., Gavryushkina A., Heled J., Jones G., Kühnert D., De Maio N., Matschiner M., Mendes F.K., Müller N.F., Ogilvie H.A., du Plessis L., Popinga A., Rambaut A., Rasmussen D., Siveroni I., Suchard M.A., Wu C-H., Xie D., Zhang C., Stadler T., Drummond A.J. 2019. BEAST 2.5: an advanced software platform for Bayesian evolutionary analysis. PLOS Comput. Biol. 15:e1006650.

12. Brodkorb P. 1972. Neogene fossil jays from the Great Plains. Condor. 74:347–349.

13. Brown J.W., Rest J.S., García-Moreno J., Sorenson M.D., Mindell D.P. 2008. Strong mitochondrial DNA support for a Cretaceous origin of modern avian lineages. BMC Biol. 6:6.

14. Cenizo M.M., Tambussi C.P., Montalvo C.I. 2012. Late Miocene continental birds from the Cerro Azul Formation in the Pampean region (central-southern Argentina). Alcheringa. 36:47–68.

15. Crouch N.M.A., Tobias J.A. 2022. The causes and ecological context of rapid morphological evolution in birds. Ecol. Lett. 25:611–623.

16. dos Reis M., Yang Z. 2013. The unbearable uncertainty of Bayesian divergence time estimation. J Syst. Evol. 51:30–43.

17. Drummond A.J., Ho S.Y.W., Phillips M.J., Rambaut A. 2006. Relaxed phylogenetics and dating with confidence. PLOS Biol. 4:e88.

18. Duchêne D., Chowdhury A.-A., Yang J., Iglesias-Carrasco M., Stiller J., Feng S., Bhatt S., Gilbert M.T.P., Zhang G., Tobias J.A., Ho S.Y.W. 2025. Drivers of avian genomic change revealed by evolutionary rate decomposition. Nature in press.

19. Duchêne S., Lanfear R., Ho S.Y.W. 2014. The impact of calibration and clock-model choice on molecular estimates of divergence times. Mol. Phylogenet. Evol. 78:277–289.

20. Ericson P.G.P., Johansson U.S. 2003. Phylogeny of Passerida (Aves: Passeriformes) based on nuclear and mitochondrial sequence data. Mol. Phylogenet. Evol. 29:126–138.

21. Ericson P.G.P., Klopfstein S., Irestedt M., Nguyen J.M.T., Nylander J.A.A. 2014. Dating the diversification of the major lineages of Passeriformes (Aves). BMC Evol. Biol. 14:8.

22. Gandolfo M.A., Nixon K.C., Crepet W.L. 2008. Selection of fossils for calibration of molecular dating models. Ann. Mo. Bot. Gard. 95:34–42.

23. Gavryushkina A., Heath T.A., Ksepka D.T., Stadler T., Welch D., Drummond A.J. 2017. Bayesian total-evidence dating reveals the recent crown radiation of penguins. Syst. Biol. 66:57–73.

24. Gavryushkina A., Zhang C. 2020. Total-evidence dating and the fossilized birth-death model. In: The Molecular Evolutionary Clock: Theory and Practice (ed. Ho S.Y.W.). Springer, Cham. pp. 175–193.

25. Gibb G.C., England R., Hartig G., McLenachan P.A., Taylor Smith B.L., McComish B.J., Cooper A., Penny D. 2015. New Zealand passerines help clarify the diversification of major songbird lineages during the Oligocene. Genome Biol. Evol. 7:2983–2995.

26. Gill F., Donsker D., Rasmussen P. (eds) (2024). IOC World Bird List (v14.2). DOI 10.14344/IOC.ML.14.2.

27. Goloboff P.A., Pittman M., Pol D., Xu X. 2019. Morphological data sets fit a common mechanism much more poorly than DNA sequences and call into question the Mkv model. Syst. Biol. 68:494–504.

28. Heath T.A., Huelsenbeck J.P., Stadler T. 2014. The fossilized birth-death process for coherent calibration of divergence-time estimates. Proc. Natl. Acad. Sci. USA 111:E2957–E2966.

29. Ho S.Y.W., Phillips M.J. 2009. Accounting for calibration uncertainty in phylogenetic estimation of evolutionary divergence times. Syst. Biol. 58:367–380.

30. Höhna S., Stadler T., Ronquist F., Britton T. 2011. Inferring speciation and extinction rates under different sampling schemes. Mol. Biol. Evol. 28:2577–2589.

31. Jarvis E.D., Mirarab S., Aberer A.J., Li B., Houde P., Li C., Ho S.Y.W., Faircloth B.C., Nabholz B., Howard J.T., Suh A., Weber C.C., da Fonseca R.R., Li J., Zhang F., Li H., Zhou L., Narula N., Liu L., Ganapathy G., Boussau B., Bayzid M.S., Zavidovych V., Subramanian S., Gabaldón T., Capella-Gutiérrez S., Huerta-Cepas J., Rekepalli B., Munch K., Schierup M., Lindow B., Warren W.C., Ray D., Green R.E., Bruford M.W., Zhan X., Dixon A., Li S., Li N., Huang Y., Derryberry E.P., Bertelsen M.F., Sheldon F.H., Brumfield R.T., Mello C.V., Lovell P.V., Wirthlin M., Schneider M.P., Prosdocimi F., Samaniego J.A., Vargas Velazquez A.M., Alfaro-Núñez A., Campos P.F., Petersen B., Sicheritz-Ponten T., Pas A., Bailey T., Scofield P., Bunce M., Lambert D.M., Zhou Q., Perelman P., Driskell A.C., Shapiro B., Xiong Z., Zeng Y., Liu S., Li Z., Liu B., Wu K., Xiao J., Yinqi X., Zheng Q., Zhang Y., Yang H., Wang J., Smeds L., Rheindt F.E., Braun M., Fjeldsa J., Orlando L., Barker F.K., Jønsson K.A., Johnson W., Koepfli K.P., O’Brien S., Haussler D., Ryder O.A., Rahbek C., Willerslev E., Graves G.R., Glenn T.C., McCormack J., Burt D., Ellegren H., Alström P., Edwards S.V., Stamatakis A., Mindell D.P., Cracraft J., Braun E.L., Warnow T., Wang J., Gilbert M.T., Zhang G. 2014. Whole-genome analyses resolve early branches in the tree of life of modern birds. Science 346:1320–1331.

32. Jønsson K.A., Fabre P-H., Kennedy J.D., Holt B.G., Borregaard M.K., Rahbek C., Fjeldså J. 2016. A supermatrix phylogeny of corvoid passerine birds (Aves: Corvides). Mol. Phylogenet. Evol. 94:87–94.

33. Jønsson K.A., Fabre P-H., Ricklefs R.E., Fjeldså J. 2011. Major global radiation of corvoid birds originated in the proto-Papuan archipelago. Proc. Natl. Acad. Sci. USA 108:2328–2333.

34. Kakegawa Y., Hirao K. 2003. A Miocene passeriform bird from the Iwami Formation, Tottori Group, Tottori, Japan. Bull. Natn. Sci. Mus., Tokyo, Ser C. 29:33–37.

35. Kennedy J.D., Borregaard M.K., Jønsson K.A., Marki P.Z., Fjeldså J., Rahbek C. 2016. The influence of wing morphology upon the dispersal, geographical distributions and diversification of the Corvides (Aves; Passeriformes). Proc. R. Soc. B 283:20161922.

36. Kessler J., Hir J. 2012. The avifauna in North Hungary during the Miocene. Part II. Földt. Közl. 142:149–168.

37. Lee M.S.Y., Palci A. 2015. Morphological phylogenetics in the genomic age. Curr. Biol. 25:R922–R929.

38. Lewis P.O. 2001. A likelihood approach to estimating phylogeny from discrete morphological character data. Syst. Biol. 50:913–925.

39. Lowi-Merri T.M., Gjevori M., Bocheński Z.B., Wertz K., Claramunt S. 2024. Total-evidence dating and the phylogenetic affinities of early fossil passerines. J. Syst. Palaeontol. 22:2356086

40. Luo A., Duchêne D.A., Zhang C., Zhu C-D., Ho S.Y.W. 2020. A simulation-based evaluation of tip-dating under the fossilized birth-death process. Syst. Biol. 69:325–344.

41. Luo A., Zhang C., Zhou Q., Ho S.Y.W., Zhu C-D. 2023. Impacts of taxon-sampling schemes on Bayesian tip dating under the fossilized birth-death process. Syst. Biol. 72:781–801.

42. Ma X., Zhang C., Yang L., Hedges S.B., Zhong B. 2025. New insights on angiosperm crown age based on Bayesian node dating and skyline fossilized birth-death approaches. Nat. Commun. 16:2265.

43. Manegold A. 2008a. Passerine diversity in the late Oligocene of Germany: earliest evidence for the sympatric coexistence of Suboscines and Oscines. Ibis 150:377–387.

44. Manegold A. 2008b. Earliest fossil record of the Certhioidea (treecreepers and allies) from the early Miocene of Germany. J. Ornithol. 149:223–228.

45. Manegold A. 2010. Two swallow species from the early Pliocene of Langebaanweg (South Africa). Acta Palaeontol. Pol. 55:765–768.

46. Mayr G. 2008. Phylogenetic affinities of the enigmatic avian taxon Zygodactylus based on new material from the early Oligocene of France. J. Syst. Palaeontol. 6:333–344.

47. Mayr G. 2013. The age of the crown group of passerine birds and its evolutionary significance – molecular calibrations versus the fossil record. Syst. Biodivers. 11:7–13.

48. Mayr G. 2017. New species of Primozygodactylus from Messel and the ecomorphology and evolutionary significance of early Eocene zygodactylid birds (Aves, Zygodactylidae). Hist. Biol. 29:875–884.

49. Mayr G. 2022. Paleogene Fossil Birds (Second Edition). Cham, Switzerland: Springer.

50. Mayr G., Manegold A. 2004. The oldest European fossil songbird from the early Oligocene of Germany. Naturwissenschaften. 91:173–177.

51. Mayr G., Manegold A. 2006a. New specimens of the earliest European passeriform bird. Acta Palaeontol. Pol. 51:315–323.

52. Mayr G., Manegold A. 2006b. A small suboscine-like passeriform bird from the Early Oligocene of France. Condor. 108:717–720.

53. McCullough J.M., Oliveros C.H., Benz B.W., Zenil-Ferguson R., Cracraft J., Moyle R.G., Andersen M.J. 2022. Wallacean and Melanesian islands promote higher rates of diversification within the global passerine radiation Corvides. Syst. Biol. 71:1423–1439.

54. McLoughlin S. 2001. The breakup history of Gondwana and its impact on pre-Cenozoic floristic provincialism. Austral. Syst. Bot. 49:271–300.

55. Miles D.B., Ricklefs R.E., Losos J.B. 2023. How exceptional are the classic adaptive radiations of passerine birds? Proc. Natl. Acad. Sci. USA 120:e1813976120.

56. Mourer-Chauviré C., Hugueney M., Jonet P. 1989. Découverte de Passeriformes dans l’Oligocène supérieur de France. C. R. Acad. Sci. Paris (II). 309:843–849.

57. Moyle R.G., Oliveros C.H., Andersen M.J., Hosner P.A., Benz B.W., Manthey J.D., Travers S.L., Brown R.M., Faircloth B.C. 2016. Tectonic collision and uplift of Wallacea triggered the global songbird radiation. Nat Commun 7:12709.

58. Nguyen J.M.T. 2016. Australo-Papuan treecreepers (Passeriformes: Climacteridae) and a new species of sittella (Neosittidae: Daphoenositta) from the Miocene of Australia. Palaeontol. Electron. 19.1.1A:1–13.

59. Nguyen J.M.T. 2019. A new species of bristlebird (Passeriformes, Dasyornithidae) from the early Miocene of Australia. J. Vert. Paleontol. 39:e1575838.

60. Nguyen J.M.T. 2023. The earliest record of bowerbirds (Passeriformes, Ptilonorhynchidae) from the Oligo-Miocene of northern Australia. Alcheringa. 47:475–483.

61. Nguyen J.M.T., Archer M., Hand S.J. 2018. Quail-thrush birds from the Miocene of northern Australia. Acta Palaeontol. Pol. 63:493–502.

62. Nguyen J.M.T., Hand S.J., Archer M. 2016. The late Cenozoic passerine avifauna from Rackham’s Roost Site, Riversleigh, Australia. Rec. Aust. Mus. 68:201–230.

63. Nguyen J.M.T., Worthy T.H., Boles W.E., Hand S.J., Archer M. 2013. A new cracticid (Passeriformes: Cracticidae) from the early Miocene of Australia. Emu. 113:374–382.

64. O’Reilly J.E., dos Reis M., Donoghue P.C.J. 2015. Dating tips for divergence-time estimation. Trends Genet. 31:637–650.

65. Oliveros C.H., Field D.J., Ksepka D.T., Barker F.K., Aleixo A., Andersen M.J., Alström P., Benz B.W., Braun E.L., Braun M.J., Bravo G.A., Brumfield R.T., Chesser R.T., Claramunt S., Cracraft J., Cuervo A.M., Derryberry E.P., Glenn T.C., Harvey M.G., Hosner P.A., Joseph L., Kimball R.T., Mack A.L., Miskelly C.M., Peterson A.T., Robbins M.B., Sheldon F.H., Silveira L.F., Smith B.T., White N.D., Moyle R.G., Faircloth B.C. 2019. Earth history and the passerine superradiation. Proc. Natl. Acad. Sci. USA 116:7916–7925.

66. Palastrova E.S., Zelenkov N.V. 2020. A fossil species of Eremophila and other larks (Aves, Alaudidae) from the upper Pliocene of the Selenga River Valley (Central Asia). Paleontol. J. 54:187–204.

67. Parham J.F., Donoghue P.C.J., Bell C.J., Calway T.D., Head J.J., Holroyd P.A., Inoue J.G., Irmis R.B., Joyce W.G., Ksepka D.T., Patané J.S.L., Smith N.D., Tarver J.E., van Tuinen M., Yang Z., Angielczyk K.D., Greenwood J.M., Hipsley C.A., Jacobs L., Makovicky P.J., Müller J., Smith K.T., Theodor J.M., Warnock R.C.M., Benton M.J. 2012. Best practices for justifying fossil calibrations. Syst. Biol. 61:346–359.

68. Prum R.O., Berv J.S. Dornburg A., Field D.J., Townsend J.P., Lemmon E.M., Lemmon A.R. 2015. A comprehensive phylogeny of birds (Aves) using targeted next-generation DNA sequencing. Nature 526:569–573.

69. Rambaut A., Drummond A.J., Xie D., Baele G., Suchard M.A. 2018. Posterior summarization in Bayesian phylogenetics using Tracer 1.7. Syst. Biol. 67:901–904.

70. Ramírez-Barahona S., Sauquet H., Magallón S. 2020. The delayed and geographically heterogeneous diversification of flowering plant families. Nat. Ecol. Evol. 4:1232–1238.

71. Ricklefs R.E. 2003. Global diversification rates of passerine birds. Proc. R. Soc. Lond. B 270:2285–2291.

72. Robinson D.F., Foulds L.R. 1981. Comparison of phylogenetic trees. Math. Biosci. 53:131–147.

73. Ronquist F., Klopfstein S., Vilhelmsen L., Schulmeister S., Murray D.L., Pasnitsyn A.P. 2012. A total-evidence approach to dating with fossils, applied to the early radiation of the Hymenoptera. Syst. Biol. 61:973–999.

74. Ronquist F., Lartillot N., Phillips M.J. 2016. Closing the gap between rocks and clocks using total-evidence dating. Phil. Trans. R. Soc. B 371:20150136.

75. Sansom R.S., Gabbott S.E., Purnell M.A. 2010. Non-random decay of chordate characters causes bias in fossil interpretation. Nature 463:797–800.

76. Selvatti A.P., Gonzaga L.P., de Moraes Russo C.A. 2015. A Paleogene origin for crown passerines and the diversification of the Oscines in the New World. Mol. Phylogenet. Evol. 88:1–15.

77. Shufeldt R.W. 1913. Further studies of fossil birds with descriptions of new and extinct species. B. Am. Mus. Nat. Hist. 32:285–306.

78. Stadler T. 2010. Sampling-through-time in birth-death trees. J. Theor. Biol. 267:396–404.

79. Stadler T., Kühnert D., Bonhoeffer S., Drummond A.J. 2013. Birth-death skyline plot reveals temporal changes of epidemic spread in HIV and hepatitis C virus (HCV). Proc. Natl. Acad. Sci. USA 110:228–233.

80. Steell E.M., Nguyen J.M.T., Benson R.B.J., Field D.J. 2023. Comparative anatomy of the passerine carpometacarpus helps illuminate the early fossil record of crown Passeriformes. J. Anat. 242:495–509.

81. Stervander M., Fjeldså J., Christidis L., Ericson P.G.P., Ohlson J.I., Alström P. 2020. Appendix 2: an updated chronology of passerine birds. In: Fjeldså J., Christidis L., Ericson P. G. P., (eds) The Largest avian radiation: the evolution of perching birds, or the order Passeriformes. Lynx Editions. pp 387–396.

82. Stiller J., Feng S., Chowdhury A., Rivas-González I., Duchêne D.A., Fang Q., Deng Y., Kozlov A., Stamatakis A., Claramunt S., Nguyen J.M.T., Ho S.Y.W., Faircloth B.C., Haag J., Houde P., Cracraft J., Balaban M., Mai U., Chen G., Gao R., Zhou C., Xie Y., Huang Z., Cao Z., Yan Z., Ogilvie H.A., Nakhleh L., Lindow B., Morel B., Fjeldså J., Hosner P.A., da Fonseca R.R., Petersen B., Tobias J.A., Székely T., Kennedy J.D., Reeve A.H., Liker A., Stervander M., Antunes A., Tietze D.T., Bertelsen M.F., Lei F., Rahbek C., Graves G.R., Schierup M.H., Warnow T., Braun E.L., Gilbert M.T.P., Jarvis E.D., Mirarab S., Zhang G. 2024. Complexity of avian evolution revealed by family-level genomes. Nature 629:851–860.

83. Volkova N.V., Zelenkov N.V. 2018. A scansorial passerine bird (Passeriformes, Certhioidea) from the uppermost Lower Miocene of Eastern Siberia. Paleontol. J. 52:58–65.

84. Warnock R.C.M., Parham J.F., Joyce W.G., Lyson T.R., Donoghue P.C.J. 2015. Calibration uncertainty in molecular dating analyses: there is no substitute for the prior evaluation of time priors. Proc. R. Soc. B 282:20141013.

85. Weidig I. 2010. New birds from the Lower Eocene Green River Formation, North America. In: Boles W.E., Worthy T.H., (eds) Proceedings of the VII International Meeting of the Society of Avian Paleontology and Evolution. Rec. Aust. Mus. 62:29–44.

86. Wetmore A. 1925. The systematic position of *Palaeospiza bella* Allen, with observations on other fossil birds. Bull. Mus. Comp. Zool. 67:183–193.

87. Worthy T.H., Hand S.J., Nguyen J.M.T., Tennyson A.J.D., Worthy J.P., Scofield R.P., Boles W.E., Archer M. 2010. Biogeographical and phylogenetic implications of an early Miocene wren (Aves: Passeriformes: Acanthisittidae) from New Zealand. J. Vert. Paleontol. 30:479–498.

88. Worthy T.H., Tennyson A.J.D., Jones C., McNamara J.A., Douglas B.J. 2007. Miocene waterfowl and other birds from central Otago, New Zealand. J. Syst. Palaeontol. 5:1–39.

89. Yang Z., Rannala B. 2006. Bayesian estimation of species divergence times under a molecular clock using multiple fossil calibrations with soft bounds. Mol. Biol. Evol. 23:212–226.

90. Zachos J. Pagani M., Sloan L., Thomas E., Billups K. 2001. Trends, rhythms, and aberrations in global climate change 65 Ma to present. Science 292:686–693.

91. Zelenkov N.V., Kurochkin E.N. 2012. The first representative Pliocene assemblages of passerine birds in Asia (Northern Mongolia and Russian Transbaikalia). Geobios. 45:323–334.

92. Zelenkov N.V. 2017. The revised avian fauna of Rudabanya (Hungary, Late Miocene). In: Acosta Hospitaleche C., Agnolín F.L., Haidr N., Noriega J.I., Tambussi C.P., editors. Paleontología Y Evolución de Las Aves. Contribuciones del MACN. 253–266.

93. Zhang C., Stadler T., Klopfstein S., Heath T.A., Ronquist F. 2016. Total-evidence dating under the fossilized birth-death process. Syst. Biol. 65:228–249.

